# Robust measurement of microbial reduction of graphene oxide nanoparticles using image analysis

**DOI:** 10.1101/2024.12.09.627602

**Authors:** Danielle T. Bennett, Anne S. Meyer

**Affiliations:** Department of Biology, University of Rochester, Rochester, New York, United States

**Author notes:** Address correspondence to Anne S. Meyer,.

## Abstract

*Shewanella oneidensis* (*S. oneidensis*) has the capacity to reduce electron acceptors within a medium and is thus used frequently in microbial fuel generation, pollutant breakdown, and nanoparticle fabrication. Microbial fuel setups, however, often require costly or labour-intensive components, thus making optimization of their performance onerous. For rapid optimization of setup conditions, a model reduction assay can be employed to allow simultaneous, large-scale experiments at lower cost and effort. Since *S. oneidensis* uses different extracellular electron transfer pathways depending on the electron acceptor, it is essential to use a reduction assay that mirrors the pathways employed in the microbial fuel system. For microbial fuel setups that use nanoparticles to stimulate electron transfer, reduction of graphene oxide provides a more accurate model than other commonly used assays as it is a bulk material that forms flocculates in solutions with a large ionic component. However, graphene oxide flocculates can interfere with traditional absorbance-based measurement techniques. This study introduces a novel image analysis method for quantifying graphene oxide reduction, showing improved performance and statistical accuracy over traditional methods. A comparative analysis shows that the image analysis method produces smaller errors between replicates and reveals more statistically significant differences between samples than traditional plate reader measurements under conditions causing graphene oxide flocculation. Image analysis can also detect reduction activity at earlier time points due to its utilization of larger solution volumes, enhancing color detection. These improvements in accuracy make image analysis a promising method for optimizing microbial fuel cells that use nanoparticles or bulk substrates.

**IMPORTANCE:** *Shewanella oneidensis* (*S. oneidensis*) is widely used in reduction processes such as microbial fuel generation due to its capacity to reduce electron acceptors. Often, these setups are labor intensive to operate and require days to produce results, so use of a model assay would reduce the time and expense needed for optimization. Our research developed a novel digital analysis method for analysis of graphene oxide flocculates that may be utilized as a model assay for reduction platforms featuring nanoparticles. Use of this model reduction assay will enable rapid optimization and drive improvements in the microbial fuel generation sector.

## INTRODUCTION

*Shewanella oneidensis* is a bacterium that can drive microbial fuel generation^1–5^, breakdown of azo dye pollutants^6–8^, and fabrication of nanoparticles^2^ due to its capacity for transferring electrons to electron acceptors within a medium^2,7,9–11^. Natively, this electron transferral facilitates bacterial respiration using extracellular electron acceptors like Fe(III)^12^. Electron transfer by the *S. oneidensis* organism is accomplished through several alternative routes intercellularly and extracellularly, including electron shuttling by flavin secretion^13–16^, through outer membrane extensions produced by the bacterium^10,17,18^, or through direct membrane pathways^19,20^. Flavins are soluble shuttle molecules which are released by the bacteria into solution^21^. In both the nanowire and the direct membrane pathways, metal reduction has been largely attributed to a set of surface Mtr/Omc proteins^22,23^ that includes three outer membrane decaheme c-type cytochromes (MtrC, MtrF, OmcA), two periplasmic decaheme cytochromes (MtrA and MtrD), and two outer membrane non-cytochrome proteins (MtrB, MtrE)^22–26^. These proteins funnel electrons to cytochromes embedded in the outer membrane of the bacterium, allowing extracellular electron transfer in particular for metallic iron electron acceptors^3,24,27^. This capacity also enables use of this organism to reduce a wide range of extracellular electron acceptors^11^, including desalination of salt water^6–8,28^, bioremediation of pollutants in soil^28^ and powering environmental sensors^4,29^.

Since microbial fuel cell setups can be complex and require extended time periods to produce results^4,28^, efficiency of the fuel cells may be improved using a model reduction assay to quickly test new reaction conditions. The reduction of the acceptor in microbial fuel cells may proceed through any of the *S. oneidensis* electron transfer pathways, and the specific reduction pathway that is utilized by the bacterium varies in different circumstances^16,30,31^. Flavin pathways seem to dominate in systems with insoluble electron acceptors and under certain ratios of electron donors and acceptors^16,30,31^, nanowire pathways seem to dominate when the organism is under stress due to nutrient deficiency^3,10,18,32^, and Mtr pathways are important in anaerobic conditions^33–35^. The dominant pathway can also depend on the type of electron acceptor present in the extracellular environment^21,36,37^. Thus, to improve the efficiency of microbial fuel-generating setups utilizing a model reduction assay, the model reduction assay must utilize the same reduction pathway as the fuel cell.

Model assays to monitor *S. oneidensis* reduction frequently involve observing the variation in absorbance over time as dye compounds such as methylene blue^13,38^ or Congo red^13^ are reduced and become colorless. Another common assay, the ferrozine assay, creates conditions of excess iron under which an iron transport pathway termed FoeE is particularly important in reduction^39^. The ferrozine assay is representative of reduction pathways involving soluble metal ions in solution or media with excess iron^39^, while Congo red reduction features soluble electron acceptors, particularly those which utilize the Mtr pathway with dependence on the CymA cytochrome^13^. For insoluble electron acceptors such as nanoparticles or bulk conductive solids, these assays will not be appropriately representative. Plasmonic nanoparticles may also be used to monitor microbial electron transfer as observed by Graham et al. (2022)^40^, and this approach offers direct insight into nanoparticle pathways and direct monitoring of electron transfer. However, this method is similar to electrochemical monitoring of microbial electron transfer in that it offers precise quantification of microbial electroactivity at small scale^41–44^. For rapid tuning of microbial fuel setups, which are usually performed at bulk scale, a graphene oxide reduction assay is advantageous as graphene oxide is inexpensive, scalable, and can be reduced rapidly^45^. Graphene oxide forms insoluble nanoparticle flocculates in solution with sizes ranging from 1 nm in diameter in colloidal solution to 20 µm in acidic or high-concentration conditions^46,47^. Graphene oxide reduction by *S. oneidensis* has been shown to require the Mtr respiratory pathway^15^ with particular dependence on MtrF and CymA^48^. This reduction process is also enhanced by electron shuttling pathways, which are more prominent when the electron acceptor is a bulk substrate^15^. Graphene oxide reduction reactions can be performed under conditions that cause the graphene oxide to flocculate, a type of aggregation, which can take place in solution under certain pH and concentration conditions^46^. *S. oneidensis* reduction of graphene oxide flocculates may thus serve as a model assay for systems that reduce nanoparticles or bulk substrate. Graphene oxide flocculates form rapidly^46,49^ and do not require the technical complexity of nanoparticle synthesis^23^, making them well-suited for use as a model system of a microbial fuel cell that features nanoparticles. However, monitoring the absorbance of graphene oxide using traditional spectroscopic methods is challenging due to this flocculation, which causes increased error in readings due to drift of the large particles. Typically, assays are restricted to certain pH ranges and graphene oxide concentrations^45^, or alternatively flocculates can be sonicated and filtered to remove them prior to analysis^11^, both of which limit the use of these assays. Visually, reduction of graphene oxide produces a color change from light brown to dark black as the graphene oxide is reduced^23^, and this visual color difference may be used to measure directly the transfer of electrons to bulk substrate or nanoparticle electron acceptors. Hence there is an opportunity for image analysis of graphene oxide reduction to provide quantitative information on the extent of reduction of the graphene oxide over time.

Image analysis methods are commonly used in nanoparticle sensing^50,51^, in which images of the samples are analyzed for the average ‘color’ of each. Color is interpreted using a set of three vectors known as a ‘color space’^52^ such as RGB (red, green, and blue) which contains the corresponding percentage of ‘red’, ‘green,’ or ‘blue’ for each pixel. For images that include fluctuating lighting or reflections, the HSV color space (hue, saturation, and value) is more robust due to the separation of lighting information into the saturation and value vectors^52–54^. By averaging the HSV vectors of a region of interest of an image of a graphene oxide sample over time, the reduction of graphene oxide over time by *S. oneidensis* may be measured. This method is advantageous for monitoring graphene oxide solutions that have flocculates present because the averaging is performed over a larger area of the sample, thereby avoiding confounding effects from the flocculates settling within the solution. This new method of combining image analysis techniques with an unconventional graphene oxide reduction assay will enable model assays for microbial fuel cells featuring bulk substrate and nanoparticles.

## RESULTS

### Determination of the average Hue, Saturation and Value vectors for images of graphene oxide

Custom Matlab^TM^ code was written to determine the hue (H), saturation (S) and value (S) vector distribution across images of graphene oxide solutions (Figure 1A). Images of experimental samples were cropped to obtain a rectangular region of interest containing glare or reflection. Points were then selected in regions with no glare or reflections (Figure 1B). Selection of 15 points or more was found to reduce the standard error of all selected points across each image to 0.09 % or less of the vector value on average (Supplemental Figure 1). A 32 x 32 pixel box was selected around each point (Figure 1C), and the average value of each pixel within the box was calculated to obtain the average hue, saturation, and value vectors for each region of interest selected.

**Figure 1:**
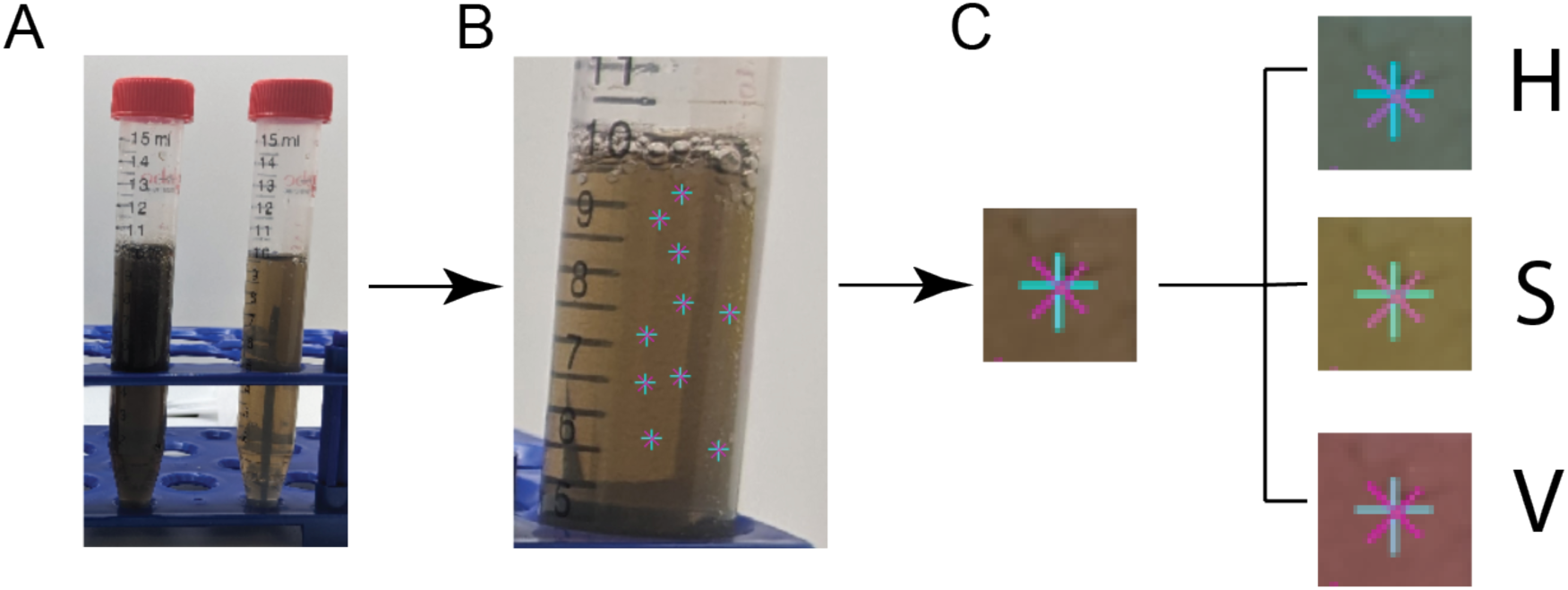
The workflow of the value vector analysis. A) A region of the image was selected that contains the least interference from the background of the sample. B) At least 12 points were selected that did not contain interference from tube labels or reflections. C) For each selected point, a 32 x 32 pixel box was cropped around the point, and the image information was separated into three vectors: hue (H), saturation (S) and value (V). The value vector contains information about the brightness or darkness of the image, represented as a number between 0 and 1, where 0 is the darkest and 1 is the lightest. The value vector was averaged over all the selected pixels to obtain an average value for each box. All boxes were then averaged to obtain an overall average value for each sample. This overall averaged value was used as a representation of the extent of graphene oxide reduction.

### Correlation between average HSV value and graphene oxide concentration

To determine the suitability of image analysis for measuring graphene oxide reduction, experiments were performed to determine whether HSV value correlated with graphene oxide concentration. Different concentrations of graphene oxide were prepared, and image analysis values for the hue, saturation, and value vectors were measured (Figure 1).

The hue and saturation vectors did not vary linearly with increasing graphene oxide concentration, rather reaching a binary saturation point (Supplemental Figure 2). However, the value vector, which is representative of the brightness of an image^54^, demonstrated linear behaviour as a function of the graphene oxide concentration in some regions of the data (Figure 2). At lower concentrations of graphene oxide, increasing the graphene oxide concentration resulted in decreasing average HSV value of the samples. The average HSV value of the samples plateaued at graphene oxide concentrations around 40 %, or 0.40 g/L. Lower concentrations of graphene oxide between 6 – 12 % were used for future experiments in order to utilize a range over which the image analysis will be sensitive. These results demonstrated that the average HSV value measurement, as opposed to the hue or saturation, showed the closest correlation with increasing graphene oxide concentration and that this measurement approach could be a promising technique for monitoring graphene oxide reduction over time.

**Figure 2:**
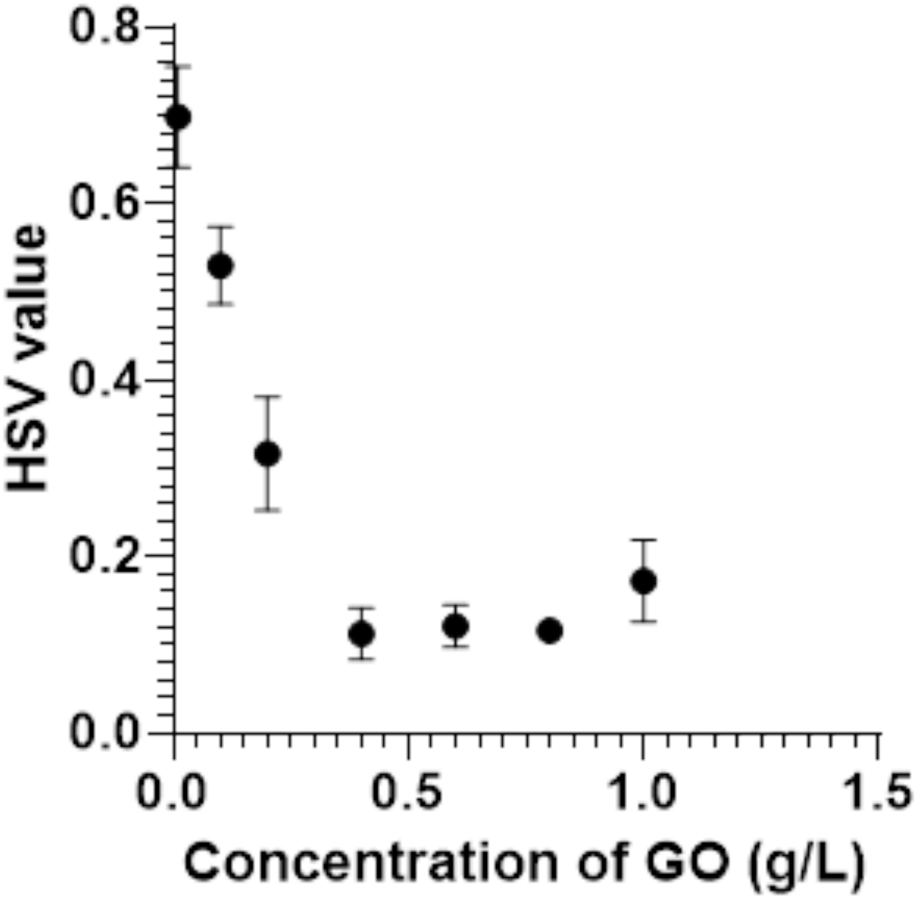
The average calculated HSV value vectors of samples with concentrations of graphene oxide varying from 0.01 – 1 g/L. Triplicate measurements were taken for each sample, and the results were averaged. The error was the standard deviation of these triplicate measurements.

### Application of the image analysis methodology by naïve users

In order to observe the user-friendliness of the image analysis technique, an experiment was performed to determine the consistency of user-selected HSV values without prior instruction. Users with no prior experience with the image analysis technique were asked to select ten or more points that accurately represented the color of an image of microbial graphene oxide reactions in 40-mL sample tubes at the 3.5-hour timepoint (Figure 3). User selections produced HSV values that were significantly statistically different, with a one-way ANOVA test resulting in a p-value of < 0.0001 (6.053, 5). Individual results for unpaired t-test p-values are presented in SI Table 1. Groups 1 and 2, 3 and 4, and 5 and 6 were not statistically different from each other but were statistically different from each other group. The average of all user-selected HSV values, considered across all users, was 0.586 ± 0.007 units, where the error represents the standard deviation of all points. This error represents 1.2% of the overall average HSV value. Hence, although different users selected statistically different values from the same sample (Supplemental Figure 3), due to the consistency of the HSV value vector across a sample the error from this variance was minimal.

**Figure 3:**
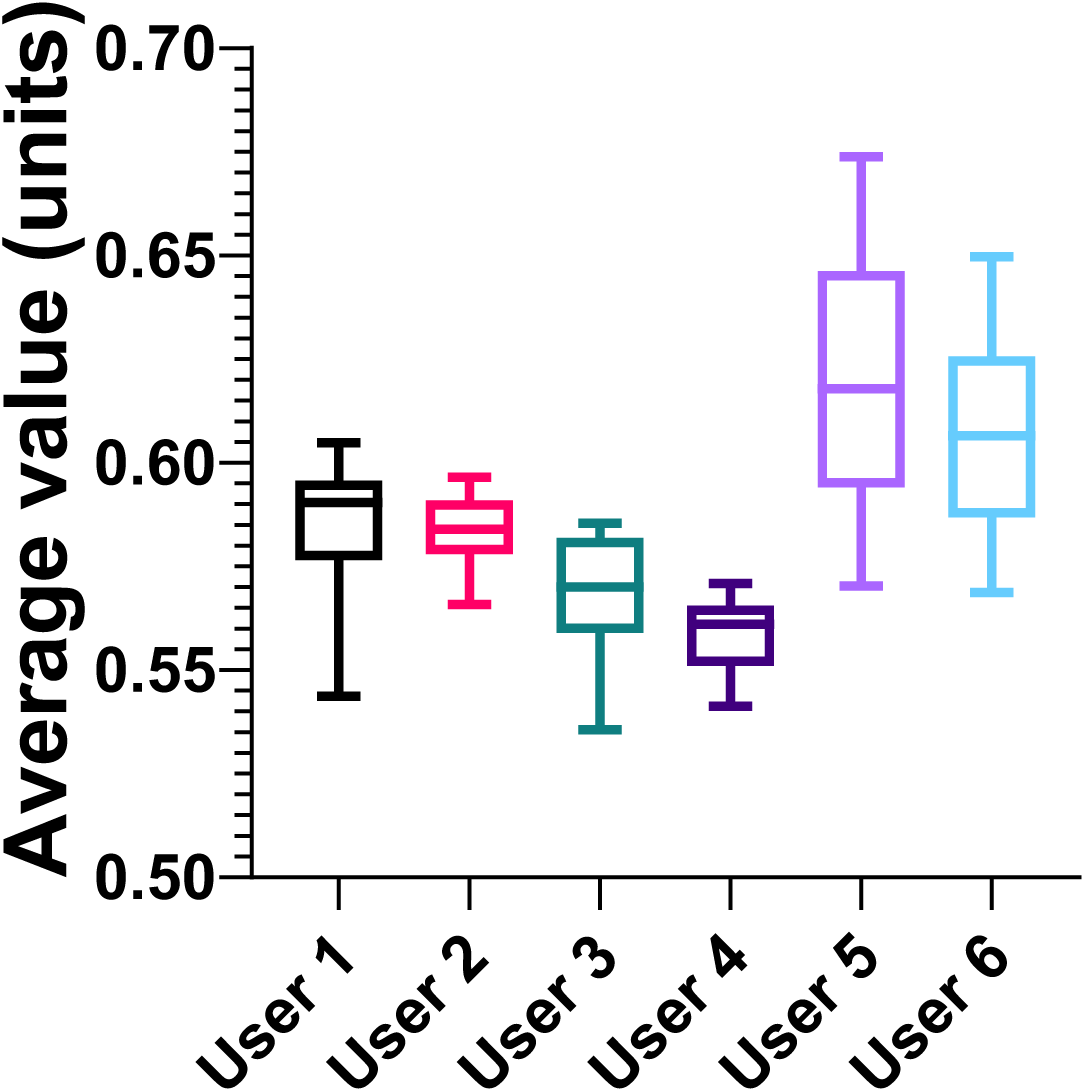
Image analysis measurements by six random users. Points were selected from the same image of three sample tubes at 3.5 hours of microbial graphene oxide reduction. The error bars represent the maximum and minimum values for each user, and the centre line represents the median. The boxes extend from the 25^th^ -75^th^ percentiles.

### Image analysis provides more sensitive measurements of graphene oxide reduction by *S. oneidensis* than plate reader absorbance measurements

Since plate reader absorbance measurements are the current standard for measuring bacterial graphene oxide reduction, experiments were performed to compare the accuracy and sensitivity of the image analysis method with plate reader absorbance measurements. A solution of graphene oxide was reduced by *S. oneidensis* bacteria either with or without lactic acid as an electron donor. For this experiment, aliquots were taken from each sample at specific time points and used for plate reader analysis (Figure 4A, Supplemental Figure 4) as well as imaging the samples, allowing for the two analysis techniques to be directly compared (Figure 4B). The lack of electron donor present in the samples with 0 mM lactic acid was expected to result in minimal bacterial reduction rates, and thus determining the level of difference between samples with and without lactic acid will allow for comparing the sensitivity of each analysis technique. For these results, initial measurements were subtracted from subsequent measurements to normalize for variation in graphene oxide amounts between samples and allow for comparison between two techniques with different y-axis units. Any negative values for the samples were assumed to be due to random fluctuation of the measurement paired with this subtraction. Plate reader absorbance measurements of timed aliquot extractions were taken at 610 nm in a 96 well plate, and at this same time-point the samples were imaged to provide a direct comparison with the image analysis.

**Figure 4:**
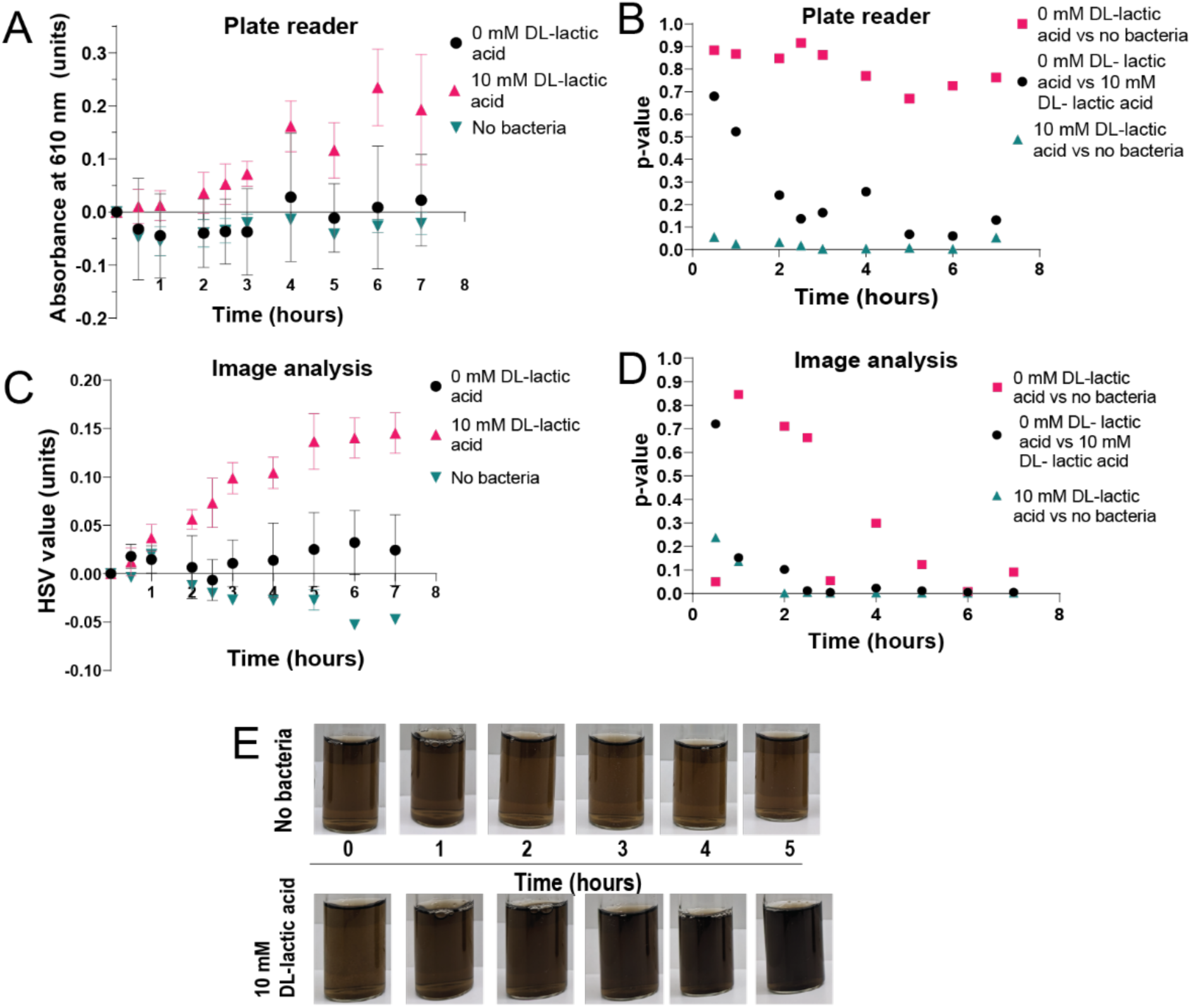
Comparison of plate reader and image analysis of bacterial graphene oxide reduction. A) Analysis of the reduction of graphene oxide by *S. oneidensis* over time by plate reader measurements of the absorbance of a 100 µL aliquot at 610 nm. The statistical difference between samples was calculated to have a p value = 0.0007, with an W ratio of 9.046 (3.000, 17.71). B) Bootstrapping statistical analysis of the p-value of different sample comparisons for the plate reader data at different timepoints. C) Image analysis measurements of the same samples, resulting in an averaged HSV value representation of the graphene oxide reduction. The statistical significance of the difference between samples was calculated to have a p-value < 0.0001 with an W ratio of 20.06 (3.000, 17.67). The errors represent the standard deviation of triplicate repeats of each sample. D) Bootstrapping statistical analysis of the p-value of different sample comparisons for the image analysis data at different timepoints. E) Images of a single sample of graphene oxide with no bacteria (top) or 10 mM lactic acid as an electron source (bottom) over time. Samples with no bacteria were used as a control to detect background levels of graphene oxide reduction. The samples with 0 mM lactic acid and with 10 mM lactic acid both contained identical initial concentrations of bacteria. The statistical significance of the difference between samples was calculated using one way ANOVA with a Welch’s correction as well as bootstrapping analysis.

The plate reader measurements showed an increase in average absorption over time after an initial lag phase for the samples containing lactic acid, indicating reduction of the graphene oxide (Figure 4A). Minimal changes in absorption were measured for the samples without lactic acid or without bacteria. The image analysis results also showed an increase in average HSV value over time for the samples containing lactic acid, though the apparent lag phase was shorter (Figure 4C). The no-lactic-acid and no-bacteria control samples again showed minimal changes in absorption. The limit of detection of a sensing assay may be estimated by obtaining the overall mean of a control sample and then adding three standard deviations to obtain an upward estimate^55,56^. To compare the different techniques, this limit of detection was normalised by the highest detected value of the assay. Since samples often had slightly different starting absorbances, the initial absorbance for an individual sample was subtracted from all subsequent values to normalize the data so that samples could be compared. The normalised value was 0.07 units for the plate reader and 0.29 units for the image analysis. This result shows that the limit of detection of the plate reader assay was lower than that of the image analysis and thus the plate reader is more sensitive. However, the lower W ratio (analogous to the F statistic^57^) for the plate reader analysis measurements indicates that the statistical difference between the samples with 10 mM lactic acid, without lactic acid, and without bacteria is lower for the plate reader measurements than for the image analysis data^57^. This result indicates that there is improved statistical separation between different sample types for the image analysis data.

To further investigate the statistical differences between sample types, bootstrapping analysis was performed on both datasets to provide an improved p-value analysis for small datasets at different time points^58^. For the plate reader analysis method (Figure 4B), the p-value did not reach a confidence interval of 95 % for the comparison of the sample with 0 mM lactic acid and the sample with no bacteria, so these groups were not significantly different. The comparison between the plate reader samples with 0 mM lactic acid and 10 mM lactic acid revealed that a 95 % confidence interval was reached after 5 hours. This result indicates that these groups became statistically significantly different at the 5-hour timepoint. The plate reader sample containing 10 mM lactic acid and the sample with no bacteria were significantly statistically different throughout the experiment. In contrast, bootstrapping analysis of the image analysis method (Figure 4D) indicated that the sample with 0 mM lactic acid and the sample with no bacteria reached statistically significant separation after 6 hours, and the 0 mM lactic acid sample and the 10 mM lactic acid sample reached statistically significant separation earlier at 2.25 hours. The sample with 10 mM lactic acid and the control sample with no bacteria became significantly statistically different at 2 hours. Since the three groups were expected to have a large difference in the rate of extracellular reduction for graphene oxide, reaching significant statistical separation at earlier timepoints indicates improved precision of the image analysis technique.

Visual comparison of the vials of two different samples over time, with one containing no bacteria and one with 10 mM lactic acid added as an electron donor, showed a qualitative difference at the two-hour time point, with the 10 mM lactic acid sample appearing darker (Figure 4E). This difference was reflected in the image analysis data but not the plate reader measurements. Reduction was indicated by the separation between the control and samples with lactic acid. The image analysis data trendline (Figure 4C) also plateaus around the five-hour time point, whereas the plate reader data does not show a clear plateau (Figure 4A). For bacterial reduction, a Gompertz trendline exhibiting a lag period, a period of exponential growth and then a plateau is expected^59,60^. This fit is often used for bacterial growth curve fitting^59,61^ and is applicable for bacterial consumption of analytes^59^. The plateau in the image analysis data trendline (Figure 4C) could indicate expected Gompertz reduction activity, or it could indicate that the sensor was saturated. The clear plateau for the image analysis method that is not present for the plate reader analysis method is indicative of the higher precision of the image analysis technique.

### Influence of differing initial bacterial densities on measurements of graphene oxide reduction by *S. oneidensis*

Since bacterial cells^62^ and reduced graphene oxide^63^ can have overlapping optical density absorbance peaks, techniques to measure graphene oxide reduction can potentially be confounded by the density of bacteria cells in the samples. Experiments were thus performed to examine the impact of differing initial O.D._600_ densities of bacteria on both the plate reader and image analysis techniques for measuring bacteria graphene oxide reduction. To avoid a plateau from analyte consumption and improve bacterial growth, electron donors were provided in excess via lysogeny broth. Providing too much excess lactic acid in combination with the high ion concentrations of the Minimal media causes near-instantaneous flocculation of graphene oxide into large flakes that settle too fast for imaging^46^. To circumvent this issue, lysogeny broth was used to provide electron donors. *S. oneidensis* bacteria were added at initial O.D._600_ densities ranging from 0.02 to 0.3. Timepoint aliquots were analysed by the plate reader and image analysis for direct comparison (Figure 5, Supplemental Figure 5) as outlined previously.

**Figure 5:**
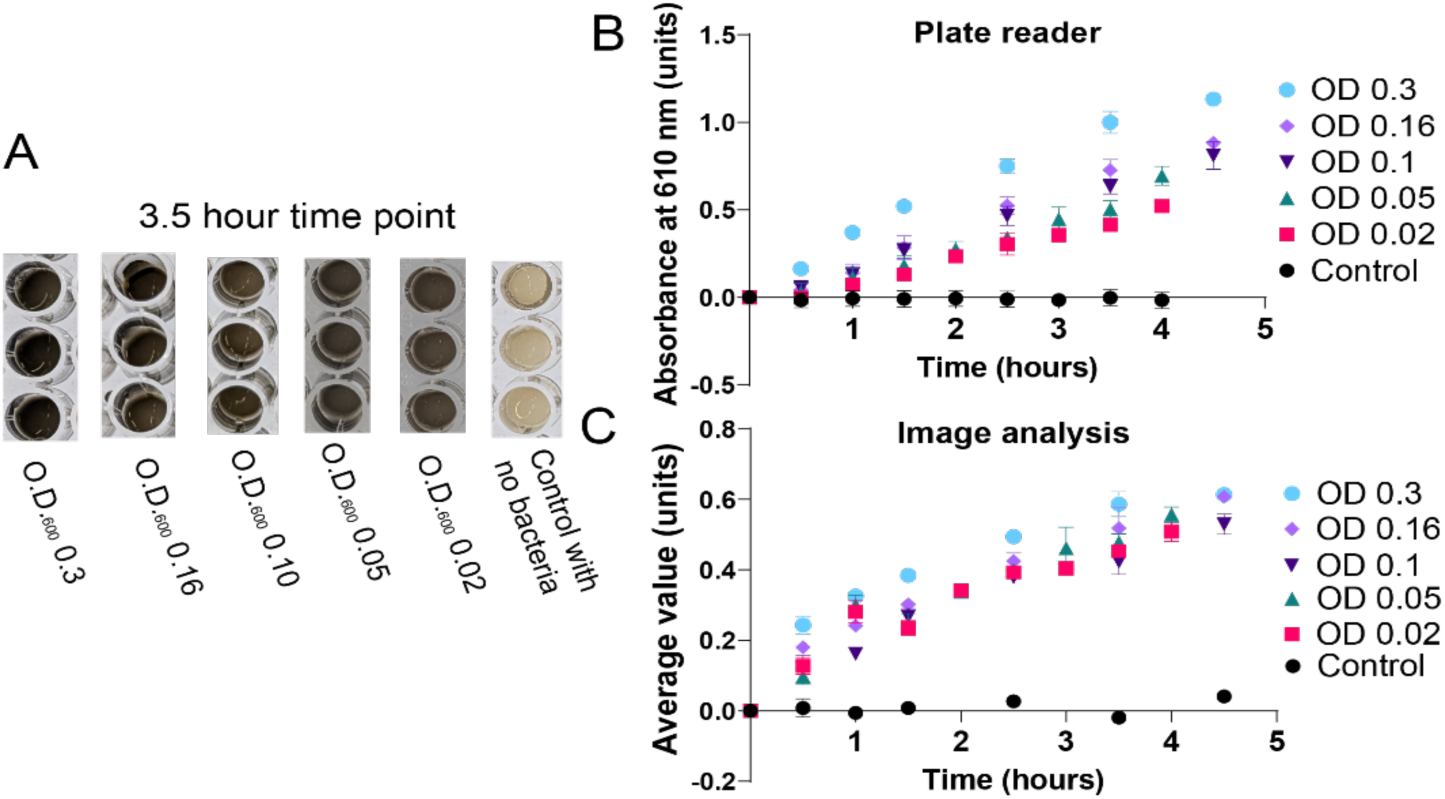
Comparison of plate reader and image analysis of bacterial graphene oxide reduction for differing initial bacterial densities. A) An example image of microplate wells containing graphene oxide solutions with different initial O.D._600_ concentrations of *S. oneidensis* at the 3.5-hour timepoint. B) Plate reader and C) image analysis measurements of these samples over time. Each sample was measured in triplicate, and the errors are representative of the standard deviation of these triplicate measurements. Some errors were too small to visualize on the graph. The control samples contained no bacteria. The statistical significance of the difference between sample sets was calculated using one way ANOVA. For comparison of samples with the control sample, the F statistic for the plate reader data was 5.158 (5, 42) with a p value = 0.0009. The F statistic for the image analysis data was 2.131 (5,40), with a p value = 0.0814.

Observation of the samples at a timepoint of 3.5 hours (Figure 5A) demonstrated dramatic differences between the control sample and the bacteria-containing samples, which were much darker in appearance. A one-way ANOVA analysis of the plate reader measurement data (Figure 5B) resulted in a p-value of 0.0009 (5.158, 5), showing significant statistical differences between groups, while the image analysis data (Figure 5C) resulted in a p-value of 0.0814 (2.131,5), showing non-significant statistical differences between groups overall. A full comparison of unpaired t-test p-values for the plate reader analysis and the image analysis may be found in SI Tables 2-3. Comparison of individual unpaired t-test p-values revealed that both analysis methods had statistically significant differences between the control sample and the samples with bacteria. For example, the p-value of the O.D._600_ 0.3 sample compared to the control sample without bacteria was 0.0011 (4.112,14) for the plate reader analysis and 0.0007 (4.543,12) for the image analysis. Non-significant differences were observed between other samples for both analysis techniques, with p-values for the image analysis being predominantly higher on average. The exception was for the comparison between the O.D._600_ 0.3 sample and the O.D._600_ 0.02 samples, which resulted in a significant difference for the plate reader analysis, with a p-value of 0.0493 (2.153,14), and a non-significant difference for the image analysis, with a p-value of 0.4491 (0.7787,14). The plate reader measurement data (Figure 5B) showed linear trendlines for the samples containing bacteria whereas the image analysis data for the samples containing bacteria (Figure 5C) had a slight plateau. The increased difference between samples containing varying starting bacterial densities seen for the plate reader analysis may likely be attributed to differences in the background scattering of the bacteria, which increases with increasing O.D._600_ instead of increased reduction activity.

### Image analysis measurements are more accurate than plate reader measurements of graphene oxide reduction by *S. oneidensis* electroactivity-deficient strains

To determine whether the image analysis technique could be applied to *S. oneidensis* strains with varying electroactivity abilities, image analysis and plate reader analysis observations were performed for the microbial reduction of graphene oxide by electroactivity-deficient *S. oneidensis* knockout strains compared to the wild-type MR-1 strain. The knockout strains were deleted for genes affecting outer membrane chromophores (*ΔmtrDΔmtrCΔomcAΔmtrF, ΔomcAΔmtrC,* and *ΔmtrCΔomcAΔmtrF*). Since these chromophores are responsible for transmission of extracellular electrons, the electroactivity of these strains is expected to be reduced^19,64–66^. At intermediate timepoints during the reduction reactions, large differences were observed visually between the reduction level of MR-1, which showed dark coloration indicative of extensive graphene oxide reduction, and the three knockout strains and the control sample containing no bacteria, which were much lighter in color (Figure 6A, Supplemental Figure 6). Comparison of the electroactivity of the *ΔmtrDΔmtrCΔomcAΔmtrF* strain with four electroactivity genes knocked out and the *ΔmtrCΔomcAΔmtrF* strain with three genes knocked out using an unpaired t-test produced a p-value of 0.6922 (0.4404, 26) for the plate reader analysis (Figure 6B) and 0.1712 (1.407, 26) for the image analysis (Figure 6C), indicating that there is not significant difference in electroactivity between the two strains. Statistical comparison of the results for the triple knockout *ΔmtrCΔomcAΔmtrF* strain versus the double knockout *ΔomcAΔmtrC* strain revealed a p-value of 0.3478 (0.9563, 26) for the plate reader analysis (Figure 6B) and 0.0160 (2.573, 26) for the image analysis (Figure 6C), indicating that only the image analysis detected a significant difference between the two strains. Comparison between the MR-1 wild-type strain and the quadruple knockout *ΔmtrDΔmtrCΔomcAΔmtrF* strain indicated a p-value of 0.0568 (1.994, 26) for the plate reader analysis (Figure 6B) and 0.0004 (4.024,26) for the image analysis (Figure 6C). The dramatic visual differences between the wild-type and knockout strains are in agreement with the image analysis results, and also align with previous observations of major differences in electroactivity between the strains^19,64–66^, indicating that the image analysis measurements are a more accurate representation of the reduction reaction in these electroactivity-deficient strains.

**Figure 6:**
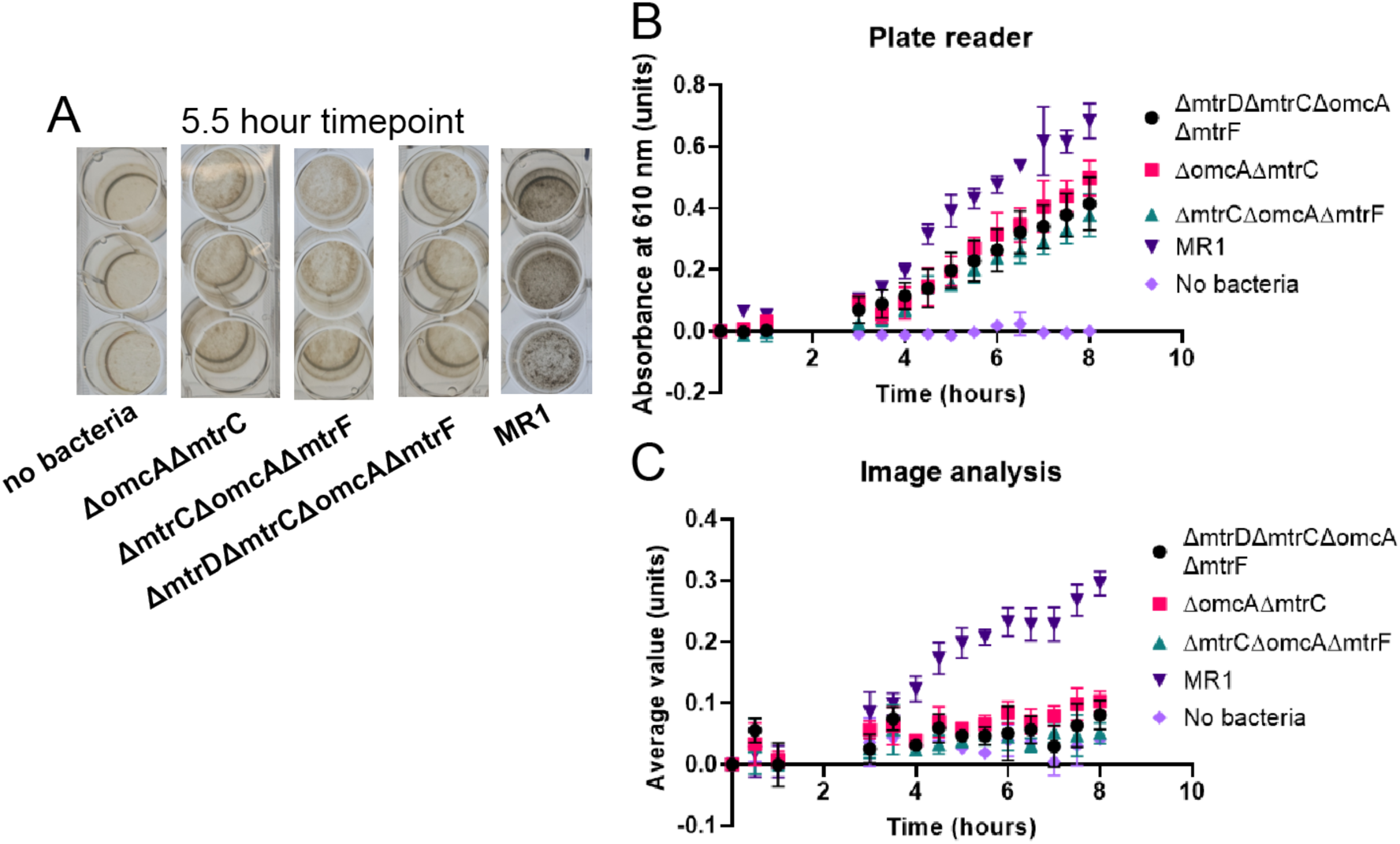
Comparison of plate reader and image analysis measurements on the electroactivity of *S. oneidensis* electroactivity-deficient strains. A) Image of plate reader wells at the 5.5-hour timepoint for graphene oxide reduction reactions containing no bacteria, *S. oneidensis* knockout strains *(ΔomcAΔmtrC, ΔmtrCΔomcAΔmtrF,* or *ΔmtrDΔmtrCΔomcAΔmtrF)*, or wild-type *S. oneidensis* MR-1. B) Plate reader analysis results for three replicate samples of each strain. C) Image analysis results for six replicate samples of each strain. Errors are standard deviations of replicate samples.

### Image analysis of scaled-up graphene oxide reduction reactions

To determine the impact of reaction scale on image analysis of microbial graphene oxide reduction, image analyses were performed on reactions with volumes ranging from 1 mL to 200 mL. Comparison between reactions of different volumes indicated a measurable impact of the scale of the reaction on the reduction rate (Figure 7). The 1-mL samples showed the slowest reduction over time, with a rate of 0.024 ± 0.001 units per hour. Interestingly, the 10-mL and 200-mL samples performed similarly over time, with reduction rates of 0.060 ± 0.003 units per hour and 0.063 ± 0.003 units per hour respectively. An unpaired t-test of the two samples produced a p-value of 0.8529 (0.1881, 18), indicating no significant difference. While the 40-mL sample had the highest reduction rate of 0.077 ± 0.002 units per hour, an unpaired t-test comparison between the 40-mL and the 200-mL sample produced a p-value of 0.5005, indicating no significant difference between the two groups. The higher reduction rate observed for the 40-mL samples may therefore be attributable to variable lag phases of the microbial reduction reaction among the different-volume samples. Overall, the image analysis technique was able to measure progressive microbial graphene oxide reduction over time for samples varying 200-fold in reaction volume.

**Figure 7:**
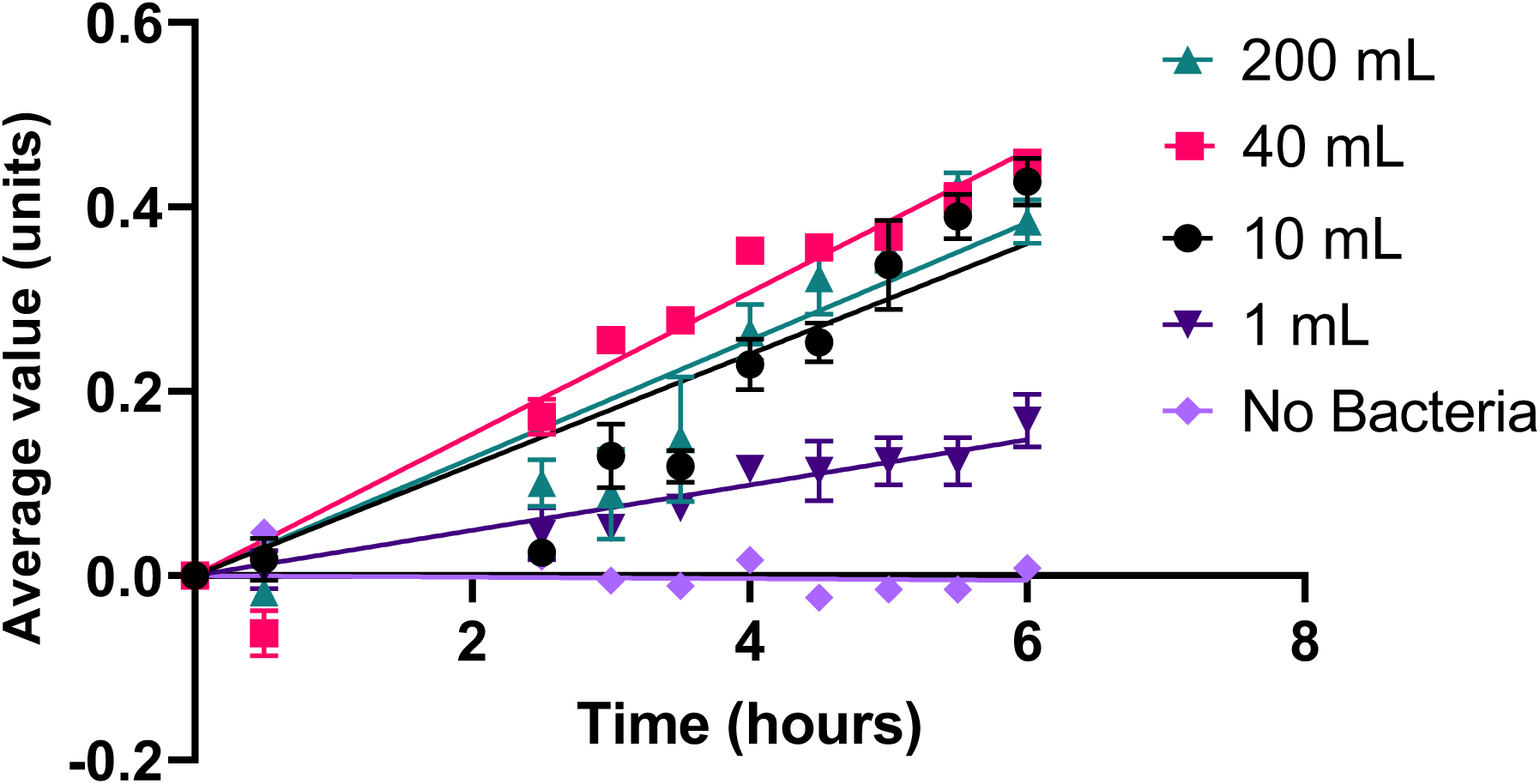
Image analysis measurements of bacterial graphene oxide reduction reactions performed with differing reaction volumes. Microbial graphene oxide reduction was performed in different volumes (200 mL, 40 mL, 10 mL, 1 mL) and monitored over time using image analysis. A control sample of 1 mL with no bacteria was included as a control. Errors are the standard deviation of three samples for each point. Linear regressions with 95 % confidence are included; regression lines pass through the origin.

## DISCUSSION

Graphene oxide reduction assays are promising candidates for application as a model assay for *S. oneidensis* reduction platforms featuring nanoparticles, since the flocculates formed by the graphene oxide would imitate nanoparticle electron acceptors^15,23^. Such a model assay would enable quick optimization of the microbial reduction platform with less resource expense. However, tailoring the electron receiver to match the original platform is important since *S. oneidensis* has multiple reduction pathways, and the dominant pathway is influenced by media composition^30^ or the specific electron acceptor^6,19,30,67^. Graphene oxide is well-suited for this purpose, but measuring the progression of the reduction reaction accurately via absorbance assays is difficult due to the flocculates. In this study, we compared traditional absorbance measurements utilizing a microwell plate reader as the gold standard^68^ with a novel digital image analysis method for a *S. oneidensis* extracellular reduction assay featuring graphene oxide flocculates^23^. The digital analysis was first optimized by determining which vector from the HSV color space would correlate most closely with graphene oxide concentration, guided by the hypothesis that correlation between an HSV vector and the graphene oxide concentration increase, which also causes similar darkening of the sample, would be representative of the color change caused by graphene oxide reduction. The HSV value vector varied linearly with increased graphene oxide within a defined range of concentrations (Figure 2), unlike the hue and saturation vectors (SI Figure 1). Hue has a sharp cut off when the colour was majority black and is thus unsuitable for monitoring gradual changes over time. Saturation was seemingly uncorrelated with the concentration of graphene oxide and was thus unsuitable. The value vector decreases with increasing graphene oxide concentration and plateaus, making it the best candidate for monitoring graphene oxide concentration using image analysis. From this data, it was hypothesized that the HSV value vector could be utilized via image analysis to be representative of graphene oxide reduction within a solution due to bacterial reduction, producing measurements varying from 0 (lightest) to 1 (darkest). A comparison of points selected by random users with no prior experience with the image analysis technique demonstrated that the HSV values of the points had an overall standard deviation between 6 different users of 1.2 % (Figure 3). This result illustrates that the image analysis technique produces consistent, reproducible results with minimal training required.

The image analysis technique was then used to measure a samples in a bacterial graphene oxide reduction assay, and these data were compared to measurements taken by a plate reader, currently the most standard method for quantifying bacterial reduction assays^68^. Analysis measurements were compared for samples containing differential levels of lactic acid as an electron donor for the *S. oneidensis.* The guiding hypothesis of this experimental set was that the samples with no electron donor would have slow extracellular reduction compared to those with electron donor present^23,69,70^ since *S. oneidensis* consumes lactic acid as an electron donor and transfers the electrons extracellularly to the graphene oxide flocculates^30,37,71^. *S. oneidensis* will eventually use the acetate it produces as a waste product as an electron source, but this reaction proceeds relatively slowly^14,72^. Thus, the statistical differences between samples would provide a measurement of the sensitivity and accuracy of the analysis method. The graphene oxide in the control solutions with no bacteria became lighter over time (Figure 4), which could be due to some larger flocculates being degraded by the shaking conditions of the experiment^73^.

Image analysis measurements (Figure 4A) showed larger differences between experimental and control samples and smaller error bars when compared to the plate reader measurements for these bacterial graphene oxide reduction experiments (Figure 4B). This increased error was likely due to two main challenges with absorbance measurements of the graphene oxide assay. Firstly, graphene oxide can form nanoparticle-like flocculates, particularly under acidic conditions^46^ and can be formed into small nanoparticles by *S. oneidensis*^23^. These flocculates settle to the bottom of the 96 microwell plate over time, which will affect the reading taken by the plate reader by variably obstructing the light source. The image analysis method can select small sections of the flakes themselves and analyse the color, and hence is not dependent on light translation through a sample. Secondly, graphene oxide absorption is usually monitored by observing the shift in the absorbance maximum from 226 nm to 262 nm for reduced graphene oxide^63^. However, due to the presence of UV-active components in the minimal media and the bacteria themselves, the intensity of this peak will be obfuscated by the other components of the media. Instead, a broad absorbance peak centred around 610 nm can be used for monitoring the reduction of graphene oxide^45^. Graphene oxide has a major absorption peak centred at 250 nm with a secondary broad peak at 450 nm^74^. As graphene oxide is reduced, the absorbance maximum becomes red-shifted, with an increase in absorbance in higher wavelengths^75^. The 610 nm absorption peak of the reduced graphene oxide overlaps with the absorption of the bacteria, usually measured at 600 nm^76^. *S. oneidensis* absorbs from 350 nm to 650 nm, exhibiting peaks at 370 nm, 540 nm and 550 nm ^77,78^ with the absorbance gradually decreasing as wavelength increases. Thus, monitoring the absorbance of GO solutions at 610 nm can detect the appearance of reduced GO without crossover with the bulk of the bacterial absorption signal. However, some interference from the bacterial scattering is unavoidable. Cheng et al. (2014)^79^ found similar interference for their electrochromic assay, which required careful monitoring of bacterial densities as analysis progressed. The image analysis does not encounter this challenge as it is not measuring absorption of a wavelength value but visual color change. This independence ensures that variations in bacterial growth do not interfere with the accuracy of measurements, which is crucial for maintaining consistency in data analysed between samples. The larger area of analysis employed for the image analysis method also reduces the impact of the flocculates. The image analysis measurements showed increased statistically significant differences between different sample types, less overlap of error bars, and the trendline contained a lag phase and a plateau consistent with the Gompertz growth expected from microbial reduction^59^ compared to the corresponding the plate reader measurements (Figure 3B). As the three samples groups are expected to have a large difference in the extracellular reduction of graphene oxide, this improved separation suggests improved sensitivity of the image analysis technique.

A plateau in graphene oxide reduction was expected to be observed at later time points of the bacteria reduction assays when the bacteria have reached high cell densities or consumed all lactate present within the sample^80^. Consumption of lactate by *S. oneidensis* has been reported at a rate of 11.8 mM/hour at high concentration of bacteria (O.D._600_ ∼ 0.2)^70^. The plateau observed by the image analysis at the five-hour time point for reduction samples containing 10 mM of lactic acid was therefore expected (Figure 3A). However, the plateau could also be due to earlier saturation of the image analysis measurement due to larger volumes of analysis. This is unlikely as further experiments obtained a Value vector maximum of 0.55 units, compared to the plateau maximum of 0.15 units from this experiment. The lack of plateau for the plate reader measurements could be due to decreased resolution of the method or increased range of measurement. Overall, the image analysis data offered high resolution compared to the greater sensitivity of the plate reader analysis method. As *S. oneidensis* is frequently used for bioreactor research, information on the rate of reduction over time is important for improving the efficiency of such systems^69,71^ so higher resolution resulting in earlier detection of reduction using the digital analysis is advantageous.

To further test the impact of bacteria concentration on graphene oxide measurements, samples containing bacteria at different initial O.D._600_ concentrations were investigated. This experiment was designed to determine whether the overlapping optical density absorbance peak of the bacteria with that of the graphene oxide would cause erroneous measurements of graphene oxide reduction in conditions with higher bacterial density. To aid the bacterial growth in this experiment, electron donors and nutrients were provided in excess. Both the plate reader analysis and image analysis showed statistically significant differences between control samples with no bacteria and samples with bacteria. These differences resulted in higher statistical difference for the image analysis. However, overall analysis of the different methods using one way ANOVA revealed larger statistical differences between groups of the plate reader analysis. Increased reduction rates with increased concentration of bacteria were not necessarily expected^81^ as *S. oneidensis* slows its electron transfer during log phase growth^82^. The differences seen for the plate reader samples, particularly those with the highest initial O.D_600_, were then likely caused by interference from the spectral overlap between the bacterial absorbance and the reduced graphene oxide absorbance measurement at 600 nm. The lack of this effect in the image analysis measurements demonstrates its effectiveness of the image analysis technique in a range of different experimental conditions, overcoming challenges presented in the absorbance measurements of graphene oxide reduction assays.

The increased accuracy of the image analysis method over plate reader measurements was also observed for reactions of graphene oxide reduction by different strains of *S. oneidensis* featuring deleted electroactivity genes (Figure 6). Larger numbers of samples could be analysed for the image analysis technique since the plate reader is only able to monitor one 24-well microplate at a time, but there is no such restriction for image analysis. Knockout strains affecting the extracellular electron transport genes *ΔmtrDΔmtrCΔomcAΔmtrF* and *ΔmtrCΔomcAΔmtrF* were revealed by unpaired t-tests to have statistically indistinguishable reduction activity for both analysis methods. However, comparison of *ΔomcAΔmtrC* and *ΔmtrCΔomcAΔmtrF* resulted in statistically significant differences only for the image analysis method and not for the plate reader analysis, as was also observed upon comparison between the MR1 wild-type strain and the *ΔmtrDΔmtrCΔomcAΔmtrF* quadruple knockout strain (Figure 6). Significant differences in graphene oxide reduction are expected between the wild-type strain and knockout strains wherein members of the extracellular electron transport pathways are deleted. For example, reduction of electroactivity in a *cymA* knockout strain of 80% was observed in comparison to the wild type^7^, and similar extreme reduction in electroactivity is consistently observed across literature^67,70,83,84^. The *ΔmtrFΔomcA* and *ΔomcAΔmtrF* strains of *S. oneidensis* were observed to reduce iron oxide at only 5 % the rate of the wild-type strain^93^. The larger differences between wild-type and knockout strains observed by the image analysis when compared to the plate reader analysis demonstrates the higher accuracy of the image analysis technique as well as its applicability to a range of microbial strains with varying electroactivity.

Another advantage of the image analysis technique when compared to plate reader analysis is that far larger solution volumes may be analysed with the image analysis. Microbial reactors are generally operated at scales of several hundred millilitres^76,85–88^, and scale can have a notable impact on catalytic reactions^89,90^. It is thus important to investigate the impact of scale upon catalytic reactions, and the use of graphene oxide reduction as a model may enable this without testing expensive catalytic components. Comparison of the microbial reduction of graphene oxide at different scales (Figure 7) demonstrates surprising results at scale. While the 1-mL samples demonstrated the slowest microbial reduction rates, the 10-mL, 40-mL, and 200-mL samples demonstrated higher reduction rates and were not significantly different from each other. These analyses indicated that the image analysis method was able to measure microbial graphene oxide reduction rates in a range of reaction volumes varying by 200-fold. The differences observed in reduction rates could derive from multiple factors that affect microbial processes at scale^91,92^. Mixing times can increase when the reaction vessel becomes larger, creating negative impacts on microbial performance due to gradients in nutrients or temperature^91^. Broth hydrostatic pressure, which rises as solution volume increases, may also disrupt microbial processes due to elevated gas partial pressure^91^. As these processes have many confounding factors, direct testing through scalable approaches such as the image analysis technique is crucial to determine the appropriateness for industrial applications^92^. Image analysis techniques are also advantageous for industrial applications due to their ease of use^79^ and the relative low cost of the technique. The smartphone used within this work cost $230 USD, in comparison to the plate reader used which cost $35,999 USD.

This work compares image analysis of bacterial graphene oxide reduction with more traditional absorption measurement collection with a microplate reader. The digital image analysis avoids increased variance from graphene oxide flocculate settling and overlap between the bacteria and the graphene oxide absorbance spectra. These features result in increased resolution of the digital image analysis, enabling earlier detection of reduction activity as well as increased statistically significant differences between different sample types. This new method creates the opportunity for a model assay for *S. oneidensis* reduction reactions that emulates interactions between the organism and nanoparticles or bulk conductive substrates. The application of this technique as a model assay could improve the speed of microbial fuel cell optimization for fuel cells containing bulk substrates or nanoparticles, accelerating the rate of advancement in this field.

## EXPERIMENTAL METHODS

### Bacterial strains and culturing

The bacterial cultures used in this work include *S. oneidensis MR-1* (ATCC® 700550™); *S. oneidensis ΔomcAΔmtrC* (JG749, Coursolle and Gralnick (2010)^93^)*; S. oneidensis ΔmtrDΔmtrCΔomcAΔmtrF* (JG1101, Coursolle and Gralnick (2010)^93^); and *S. oneidensis ΔmtrCΔomcAΔmtrF* (JG596, Coursolle and Gralnick (2010)^93^). *S. oneidensis* cultures were grown from freezer stocks stored in 25% glycerol at -80 °C. Lysogeny broth (LB) media was prepared by dissolving tryptone (0.14 M, Sigma-Aldrich), yeast extract (15 mM, Sigma-Aldrich) and NaCl (0.17 M, Sigma-Aldrich) in de-ionised water. This solution was autoclaved for 30 minutes on a fluid cycle at 121 °C and stored at room temperature. Frozen bacterial cultures were streaked onto LB-agar plates and incubated at 30 °C overnight. The following day, a single colony was isolated and grown in LB (5 mL) at 30 °C overnight under continuous shaking (200 rpm). These overnight cultures were used for graphene oxide reduction experiments.

### Graphene oxide reduction assay

A solution of Minimal media was prepared as in Edwards, Jelusic, Kosko, McClelland, Ngarnim, Chiang, Lampa-Pastirk, Krauss, and Bren (2023)^94^; details are presented in SI Tables 4-7. To this solution, graphene oxide (0.12 g / L) was added. This solution was divided into 20 mL aliquots in individual 40 mL Environmental Protection Agency (EPA) vials (Sigma-Aldrich 23188). DL-lactic acid (10 mM, lactic acid) was added to some samples as an electron donor^37^. All samples were adjusted to pH 7 with 10 M sodium hydroxide. Nitrogen gas was bubbled though each vial for 10 minutes to create anaerobic conditions, which promotes increased lactate consumption by the *S. oneidensis*^30^. Overnight cultures of *S. oneidensis* prepared as described above were added to each vial at a 1:100 dilution, with three vials lacking bacteria to serve as a negative control. The samples were incubated at 30 °C under continuous shaking (200 rpm).

### Graphene oxide reduction assay comparing different bacteria optical densities

A frozen culture of *S. oneidensis* was streaked onto an LB-agar plate then grown overnight at 30 °C. A single colony was isolated and grown in LB media at 30 °C under continuous shaking (200 rpm). The optical density (O.D._600_) of the culture was measured hourly using a Thermo Fisher Scientific™ Nanodrop to observe the absorbance of the solution at a wavelength of 600 nm. When the culture reached exponential growth phase (at an O.D._600_ of 0.5), the bacterial culture was diluted to achieve variable O.D._600_ values and was added to graphene oxide (0.12 g/L) in LB that had been flushed with nitrogen for 10 minutes to achieve anaerobic conditions.

### Generation of plate reader and image data

At specific time points, the samples prepared above were placed in a light box to reduce interference from changing lighting and background and imaged with a Google© Pixel phone. RAW image files were used for analysis to avoid any interference from the imaging software. RAW image files were converted to JPEG files using Adobe Lightroom© with no editing. The JPEG images were used for image analysis in MATLAB©.

Immediately after imaging, 100 µL aliquots were taken from each sample. The aliquots were placed into a 96 well plate, and their optical absorbance at 610 nm was measured using a Biotek© Synergy H1 microplate reader. These concurrent measurements enabled direct comparison between the measurements of graphene oxide reduction by digital image analysis and plate reader absorbance measurements of the same samples.

### Color analysis of graphene oxide reduction sample images

The JPEG images obtained of graphene oxide reduction samples were subjected to further image analysis to obtain the average of the HSV value vector of each sample at different time-points. A workflow of the analysis procedure is presented in Figure 1. Analysis code was written in-house in MATLAB© and is available in the SI. For each image, the sample was selected and cropped to select an area that had minimal interference from background or reflections (Figure 1A). The orientation, location, focus, and selected area were kept consistent between images within an experiment. Within the selected area, at least 12 points were selected, avoiding any regions featuring reflections (Figure 1B). Around each point, a 32 x 32 pixel box was generated. The color data was represented by the HSV color space, which contains the color information of the selected crop box with three vectors^52^ (Figure 1C). The H vector contains hue information: the color of each pixel. The S vector contains saturation information: how intense each pixel is. The V vector contains value information: how light or dark each pixel is. The HSV color space is advantageous over other color spaces for image analysis, such as the more typical RGB, because the data obtained is more stable under changing lighting conditions^53^. The value vector was isolated, and the average value was taken over the entire box. Each pixel was represented by a number between 0 and 1, where 0 is the darkest and 1 is the lightest. This average value was calculated for each selected point, and the average overall value across all the points was determined to obtain an average across each sample. This method allowed for determination of the standard deviation over the sample for the analyzed points.

### Image analysis of different graphene oxide concentrations

To determine the correlation of the value vector with the concentration of graphene oxide, different percentages of graphene oxide were mixed with LB media. Graphene oxide percentages ranged from 0.1% - 100%. Images were taken of the different samples in a lightbox, and the images were analysed digitally as described in the previous section. Average HSV vectors of the samples were compared to observe whether higher concentrations of graphene oxide corresponded to higher vector measurements.

### Analysis of graphene oxide reduction by *S. oneidensis* electroactivity-deficient strains

Three identical 24-well tissue culture plates (Fisherbrand, Thermo-Fisher Scientific) were prepared, each containing 3 replicate samples of *S. oneidensis* knockout strains *ΔmtrDΔmtrCΔomcAΔmtrF, ΔomcAΔmtrC, ΔmtrCΔomcAΔmtrF*, or wild-type strain MR-1. Each sample well contained 1 mL LB media with 5% (0.05 g) graphene oxide and was inoculated with a 1:100 dilution of overnight bacteria culture. Samples were sealed with optically clear tape (McMaster-Carr) to prevent evaporation and were incubated at 30°C. Two of these plates were analysed using image analysis, and one plate was analysed using the Biotek© Synergy H1 microplate reader with measurements taken at 610 nm absorbance for 26 hours.

### Image analysis of graphene oxide reduction at different scales

An overnight culture of MR-1 was inoculated at 1:100 dilution into LB media containing 5% graphene oxide (1 g / L), using volumes of 1 mL, 10 mL, 40 mL, or 200 mL. The solutions were incubated at 30 °C in 24 well tissue culture plates (Fisherbrand, Thermo-Fisher Scientific), 40 mL EPA vials (Sigma-Aldrich), 50 mL centrifuge tubes (Sigma-Aldrich), or 250 mL media bottles (Sigma-Aldrich) respectively. Imaging was performed using a Miroco SAD lamp (Miroco) in a lightbox for even illumination. Error was calculated as the standard deviation of three replicate samples.

### Application of the image analysis methodology by different users

Six different users with no prior exposure to the image analysis methodology were asked to select a region of an image that accurately represented the color of a 40 mL centrifuge tube from the previous experiment at the 3.5-hour timepoint. Users were then asked to select ten or more points from the section they had selected that accurately represented the color. No other instruction was provided. The HSV value was then calculated for each selected point, and the median HSV value for each user’s points was determined.

### Statistical analysis of data trends

Analysis was performed using Graphpad Prism^TM^ to obtain p-values. An ANOVA test was performed with a Welch’s correction to obtain the statistical difference between different sample trends.

Bootstrapping analysis was performed following the methods of B. Efron & R. Tibshirani (1993)^58,95^. Let *x*_1_, …, *x*_*n*_ be a sample distribution from a distribution with mean *x̅* and variance σ^2^_*x*_. Let *y*_1_, …, *y_n_* be an independent sample distribution with mean *y̅* and variance σ^2^_*y*_. Then the test statistic is calculated as:

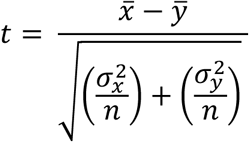

Then create two new data sets where the values are *x*^′^_*i*_ = *x_i_* − *x̅* + *z̅* and *y^′^_i_* = *y_i_* − *y̅* + *z̅*, where *z̅* is the mean of the combined sample. Draw a random sample (*x*^∗^_*i*_) of size *n* points from this pooled sample with replacement from *x*^′^_*i*_. Draw another random sample (*y*^∗^_*i*_) of size *m* points from this pooled sample with replacement from *y*^′^_*i*_. Then calculate the test statistic:

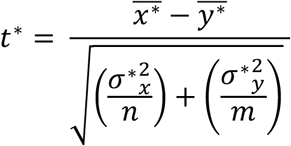

Repeat this algorithm 1,000,000 times (B = 100000) to estimate the p-value as:

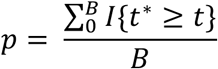

Where *I*{*t*^∗^ ≥ *t*} = 1 when the condition is true. This method is advantageous for estimating the p-value from smaller datasets^58^.

## ACKNOWLEDGEMENTS

The authors wish to thank Professor Todd Krauss and Professor Kara Bren for initial discussions concerning the spectroscopy of the graphene oxide flocculates. *Shewanella oneidensis* knockout strains were kindly gifted by Dr. Jeffrey Gralnick (University of Minnesota). The authors also wish to thank Dr. Elio Abbondanzieri for advice regarding bootstrapping analysis. The authors also wish to thank Linaria Larkin, Tiana Rohe, Jordan Kassanoff, Azra M. Walker, Dr. Elio A. Abbondanzieri, and Elliot Blues for their assistance in testing the consistency of the image analysis code. Funding to D.B. and A.S.M. was provided by the Department of Energy via DE-SC0023354.

